# Cryo-EM Structures Reveal Upstream DNA Interactions within the Mitochondrial Transcription Initiation Complex

**DOI:** 10.64898/2026.04.09.717523

**Authors:** Rory E. Sharkey, Caitlin Schroeder, Xiangyu Deng, Jamie Smith, Alfredo J. Hernandez, Yang Gao

**Affiliations:** Department of BioSciences at Rice University, Houston, TX 77005, USA; Department of Biology, Tufts University, Medford, MA 02130, USA

**Keywords:** Mitochondria, transcription, TFAM, PolRMT, transcription initiation, mitochondrial DNA (mtDNA)

## Abstract

Mitochondrial DNA (mtDNA) transcription is essential for cellular energy production and is carried out by a streamlined transcription system in which transcription factor A (TFAM), transcription factor B2 (TFB2M), and the mitochondrial RNA polymerase (PolRMT) assemble at defined promoters to initiate transcription. Previous structural studies elucidated the core initiation mechanism but relied on truncated promoter templates that excluded upstream regulatory DNA interactions. Here, we present two conformations of mitochondrial transcription initiation complexes assembled on the heavy-strand promoter (HSP): a TFAM-bound complex with extended upstream DNA and a TFAM-free complex containing short linear DNA. The TFAM-bound structure reveals a transcription-stimulatory interface between PolRMT and the upstream promoter region (UPR) enabled by TFAM-induced promoter bending. Consistent with this structural observation, UPR truncation reduces transcription from all mtDNA promoters, an effect abolished by mutation of the PolRMT interface. In contrast, the TFAM-free structure reveals a transcription-inhibitory interaction of linear upstream DNA with the PolRMT tether helix, which would sterically clash with TFAM binding. Deletion of the tether helix increases off-target transcription, supporting an autoinhibitory role that enhances promoter specificity. Together, these findings reveal how interactions of TFAM and PolRMT with upstream promoter DNA influence activity and specificity of mitochondrial transcription initiation.

## Introduction

Mitochondria orchestrate cellular life and death signals through energy production, immune signaling, and apoptotic pathways (1, 2). Central to these diverse functions is the expression of the mitochondrial genome (mtDNA), a compact and evolutionarily distinct genome housed inside the organelle (3, 4). Transcription of mtDNA results in expression of 13 core proteins involved in oxidative phosphorylation (OXPHOS), the key energy-producing pathway, as well as 22 tRNAs, 2 rRNAs, and mtDNA replication primers (3–5). As such, dysregulation of mitochondrial transcription has been linked to various disorders, including neurodegenerative and cardiovascular diseases as well as cancer (6–9).

In stark contrast to the multi-subunit, highly complex nuclear transcription machinery (10), mitochondrial transcription initiation relies on just three proteins (4, 11, 12). Transcription is carried out by PolRMT, a nuclear-encoded single-subunit RNA polymerase, structurally related to the T7 bacteriophage RNAP (13). Unlike the T7 RNAP, transcription initiation by PolRMT requires two transcription factors: transcription factor A (TFAM), which binds to and bends promoter DNA (14–16), and transcription factor B2 (TFB2M), which interacts with PolRMT near the transcription start site (TSS) to stabilize the transcription bubble (15, 17–19). Mitochondrial transcription is initiated at three promoter sequences, known as the heavy-strand promoter (HSP), the light-strand promoter (LSP), and the light-strand promoter 2 (LSP2) (20). The HSP is the principal promoter for mitochondrial gene expression as it encodes for 12 of the 13 mRNAs, 14 of the 22 tRNAs, and both of the rRNAs, while LSP and LSP2 encode the remaining RNAs (3, 4, 11). Additionally, regulation of transcription initiation is affected by TFAM-mediated compaction of mtDNA into nucleoids, where tight nucleoid compaction inhibits transcription (21). Furthermore, the non-coding 7S RNA was recently shown to stimulate formation of a transcriptionally-inactive PolRMT dimer (22). While there are far fewer components of mitochondrial transcription relative to nuclear transcription, the mechanisms and interactions influencing mitochondrial transcription initiation are nuanced and require further elucidation.

Despite extensive biochemical and structural analyses of mitochondrial transcription initiation complexes (mtTICs) (15, 23–27), critical aspects of promoter engagement remain unresolved. Notably, previously solved structures (15, 23, 24) relied on promoter templates derived from the LSP and limited to the −40 to −50 position relative to the TSS, potentially excluding upstream regulatory DNA interactions. However, DNase-footprinting and cross-linking has shown that mtTICs interact with DNA further upstream (−50 to −60) (20, 28, 29). Consistent with this observation, *in vitro* transcription using longer promoter DNA templates has demonstrated enhanced transcription activity from both LSP and HSP, with the effect being most pronounced for HSP (27, 28). Moreover, the enhanced transcription for HSP was proposed to involve binding of an additional TFAM molecule and looping of the extended DNA (28). Together, these findings suggest that intrinsic elements of the upstream DNA, particularly for the HSP, and PolRMT contribute to transcription enhancement in ways not captured by current structures.

Current structures have established interaction between TFAM and the PolRMT tether helix (15, 23, 24, 27), suggesting an essential role of TFAM in promoting PolRMT binding and mtTIC assembly. Paradoxically, although TFAM is considered essential for canonical mtTIC formation, biochemical and structural studies have demonstrated that a stable and active transcription complex can form in its absence (23, 25, 28, 30). Further, studies on truncation of the PolRMT N-terminal extension (NTE), which contains the tether helix, have shown enhanced transcription activity in Drosophila (31) and increased off-target transcription initiation of mouse PolRMT *in vitro*, suggesting an additional role of the NTE in modulating activity (25). Elucidating the interactions between TFAM, PolRMT, and promoter DNA is required to clarify how these elements influence activity and specificity of transcription initiation.

Here, we present cryo-EM analysis of an mtTIC with an extended HSP template, which revealed two distinct conformations that identify key interactions with the upstream promoter DNA. The first structure, HSP with TFAM, resolves an interaction of PolRMT with extended upstream promoter DNA, which we have termed the upstream promoter region (UPR). Consistent with the structural interface, templates containing the UPR had enhanced transcript production from all three promoters *in vitro*. Further, mutation of three key lysine residues (K425E/K428E/K432E) at the PolRMT interface ablated UPR-mediated transcription enhancement. The second conformation showed a complex with short, linear upstream DNA engaged by the PolRMT tether helix but without TFAM. Truncation of the tether helix increased transcription initiation on a template lacking the TFAM binding site, suggesting an autoinhibitory mechanism that enforces promoter specificity. Together, our study defines interactions between PolRMT and upstream DNA, which are dependent on TFAM-induced promoter bending, and reveals how these interactions modulate activity and specificity of transcription initiation.

## Results

### Cryo-EM structure of the Mitochondrial Transcription Initiation Complex on an extended Heavy-Strand Promoter

Although extended promoter regions have been shown to stimulate mitochondrial transcription initiation (28), the underlying mechanism remains unclear. Previous structural studies employed DNA substrates containing minimal promoter sequences 40 to 50-bp upstream of the TSS (15, 23, 24). To investigate the role of extended promoter DNA in mitochondrial transcription initiation, we performed single-particle cryo-EM analysis of the mtTIC with an extended HSP template. The DNA substrate consisted of the HSP sequence from −60 to +11, with a 7-bp bubble (−4 to +3) generated by mutating the non-template strand (NT), similar to the mtTIC crystal structure (15) (**Fig. 1A**). The HSP with TFAM mtTIC structure was determined at 3.33 Å resolution (**Fig. S1** and **Table 1**). We were able to resolve TFAM, TFB2M, and PolRMT along with DNA from the −57 to +11 position (**Fig. 1B**), including an extended linear DNA that approaches the N-terminal domain (NTD) of PolRMT (**Fig. 1C**).

**Figure 1.**
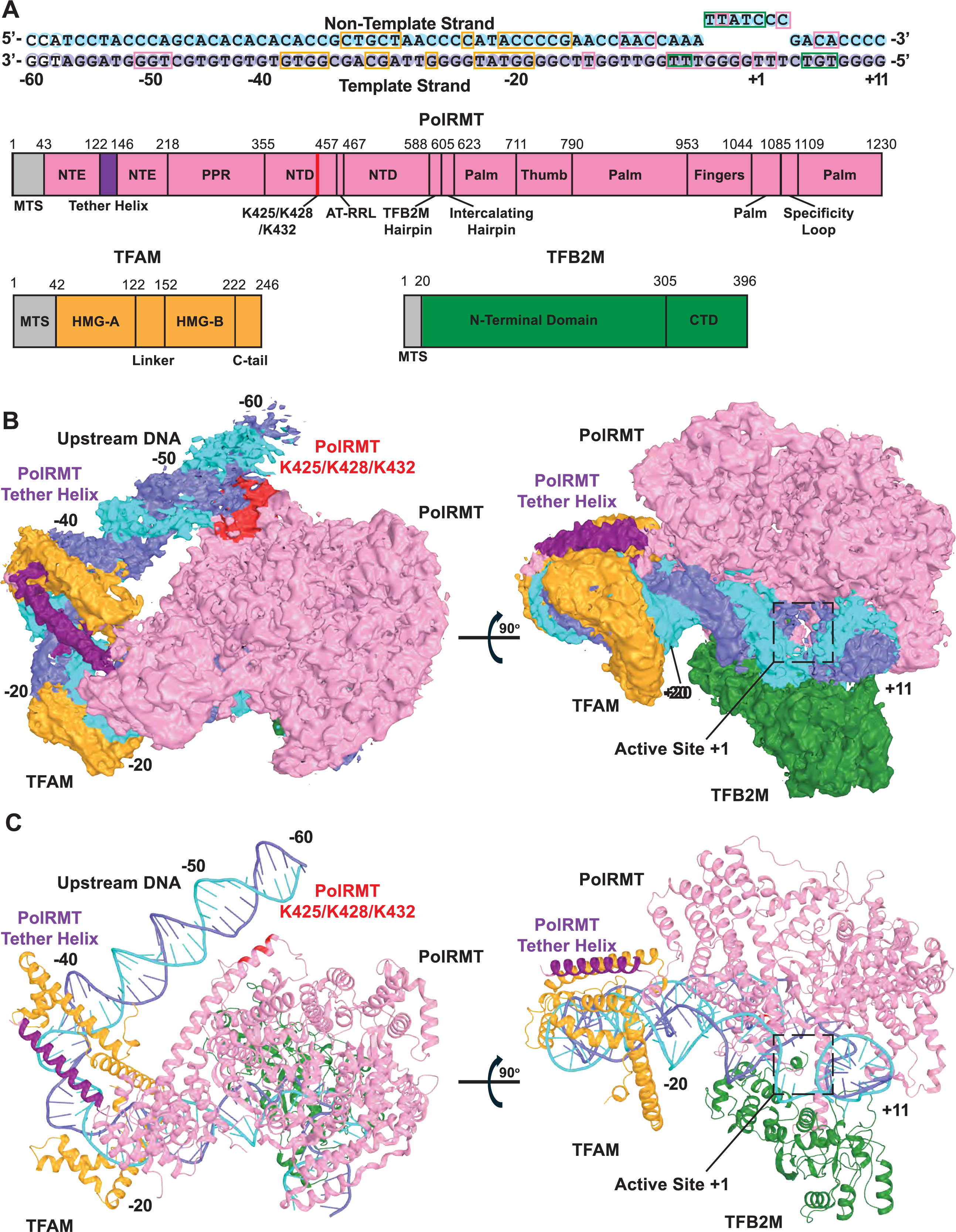
Cryo-EM structure of the Mitochondrial Transcription Initiation Complex on HSP with extended upstream DNA. (A) Schematic of the HSP bubble DNA substrate, TFAM, TFB2M, and PolRMT. The DNA substrate non-template strand (NT) and template strand (TS) are shown in cyan and slate, respectively, and shaded circles represent the resolved bases, bases that are not visible are represented by hollow circles. Protein-DNA contacts (<4 Å) are indicated by pink, green and orange boxes for PolRMT, TFB2M, and TFAM, respectively. PolRMT is shown in pink and divided into its domains: mitochondrial targeting sequence (MTS) (not included in expression constructs), N-terminal extension (NTE), tether helix in purple, pentatricopeptide repeat domain (PPR), N-terminal domain (NTD), residues K425, K428, and K432 in red, AT-rich recognition loop (AT-RRL), TFB2M-interacting hairpin (TFB2M hairpin), palm, thumb, and specificity loop. TFAM is shown in orange with the MTS, high mobility group box A (HMG-A), linker, HMG-B, and C-terminal tail (C-tail). TFB2M is shown in green with the MTS, N-terminal domain, and C-terminal domain (CTD). (B) Left: cryo-EM density map of the mtTIC on the HSP substrate showing the extended upstream DNA with proteins and DNA colored as in (A). Right: 90° rotation around the X-axis showing the PolRMT active site and downstream DNA. (C) Cartoon representation of DNA and proteins shown in (B).

**Table 1.** Cryo-EM data collection, model refinement, and validation.

|  | HSP TIC with<br>TFAM | HSP TIC without<br>TFAM |
| --- | --- | --- |
| **Data collection and processing** |  |  |
| EMDB ID |  |  |
| PDB ID |  |  |
| Microscope | Titan Krios G4i ($\Delta$) | Titan Krios G4i ($\Delta$) |
| Detector | Falcon 4i | Falcon 4i |
| Magnification | 146,000 | 146,000 |
| Voltage (kV) | 300 | 300 |
| Electron exposure ($e^-/\text{\AA}^2$) | 50 | 50 |
| Defocus range ($\mu\text{m}$) | -0.8 to -2.0 | -0.8 to -2.0 |
| Pixel size ($\text{\AA}$) | 0.958 | 0.958 |
| Number of movies | 27,688 | 27,688 |
| Initial particles (no.) | 11,635,618 | 32,940,939 |
| Final particles (no.) | 200,388 | 446,280 |
| Map resolution (Å) | 3.33 | 2.86 |
| FSC (Fourier shell correlation) threshold | 0.143 | 0.143 |
| **Refinement** |  |  |
| No. of atoms | 15733 | 12320 |
| No. of residues | - | - |
| Protein | 1606 | 1358 |
| Nucleic acid | 136 | 70 |
| RMSDs | - | - |
| Bond lengths (Å) | 0.007 | 0.007 |
| Bond angles (°) | 1.047 | 1.143 |
| **Validation** |  |  |
| MolProbity score | 1.29 | 1.39 |
| Clashscore | 5.43 | 6.99 |
| Rotamer outlier (%) | 0.00 | 0.00 |
| Ramachandran plot (%) | - | - |
| Favored | 97.99 | 98.00 |
| Allowed | 2.01 | 2.00 |
| Outliers | 0.00 | 0.00 |

The PolRMT and TFB2M core in our transcription initiation complex adopts a similar conformation as observed in previous structures (15, 23, 24). Alignment of PolRMT and TFB2M Cα atoms results in an RMSD of 1.1 Å over 1106 residues to the HSP crystal structure (PDB: 6ERQ) and 0.9 Å over 893 residues to an LSP structure solved by cryo-EM (PDB: 9MN5) (**Table S2**) (15, 23, 24). The PolRMT fingers domain adopts a clenched conformation in our structure, which is consistent with other structures that do not have an incoming NTP bound (15, 24) **(Fig. S2A**). However, TFAM is rotated by 16° in our structure compared to the HSP crystal structure (PDB: 6ERQ) and 12° to the LSP cryo-EM structure (PDB: 9MN5), suggesting flexibility of TFAM within the mtTIC (15, 23, 24) (**Fig. S2B** and **Table S2**).

### PolRMT interacts with the upstream promoter region during transcription initiation

In our mtTIC structure, clear density can be observed for the HSP UPR, which forms an additional interaction interface with PolRMT. Specifically, K425, K428, and K432, which are located in a helix of the PolRMT NTD, contact the DNA phosphate backbone through electrostatic interactions (**Fig. 2A**). While our structure reveals this interaction on the HSP, DNase footprinting experiments have suggested that the interaction occurs on all three mitochondrial promoters (20). To test the role of the UPR in transcription initiation, we designed long DNA templates including the UPR to the −70 position and truncated templates that ended at the −40 position from all three promoters (**Fig. 2B**). An *in vitro* transcription assay using the reconstituted mtTIC with PolRMT titration was employed to measure transcript production from each DNA substrate. On the LSP, the truncated template had ∼50% lower transcript production (**Fig. 2C**). Similarly, template truncation led to a decrease in transcript production for both HSP and LSP2, with a 20% and 50% reduction, respectively (**Fig. 2D-E**). Together, these results show that the PolRMT NTD interacts with the UPR and that this interaction is important for transcription initiation at all three mitochondrial promoters.

**Figure 2.**
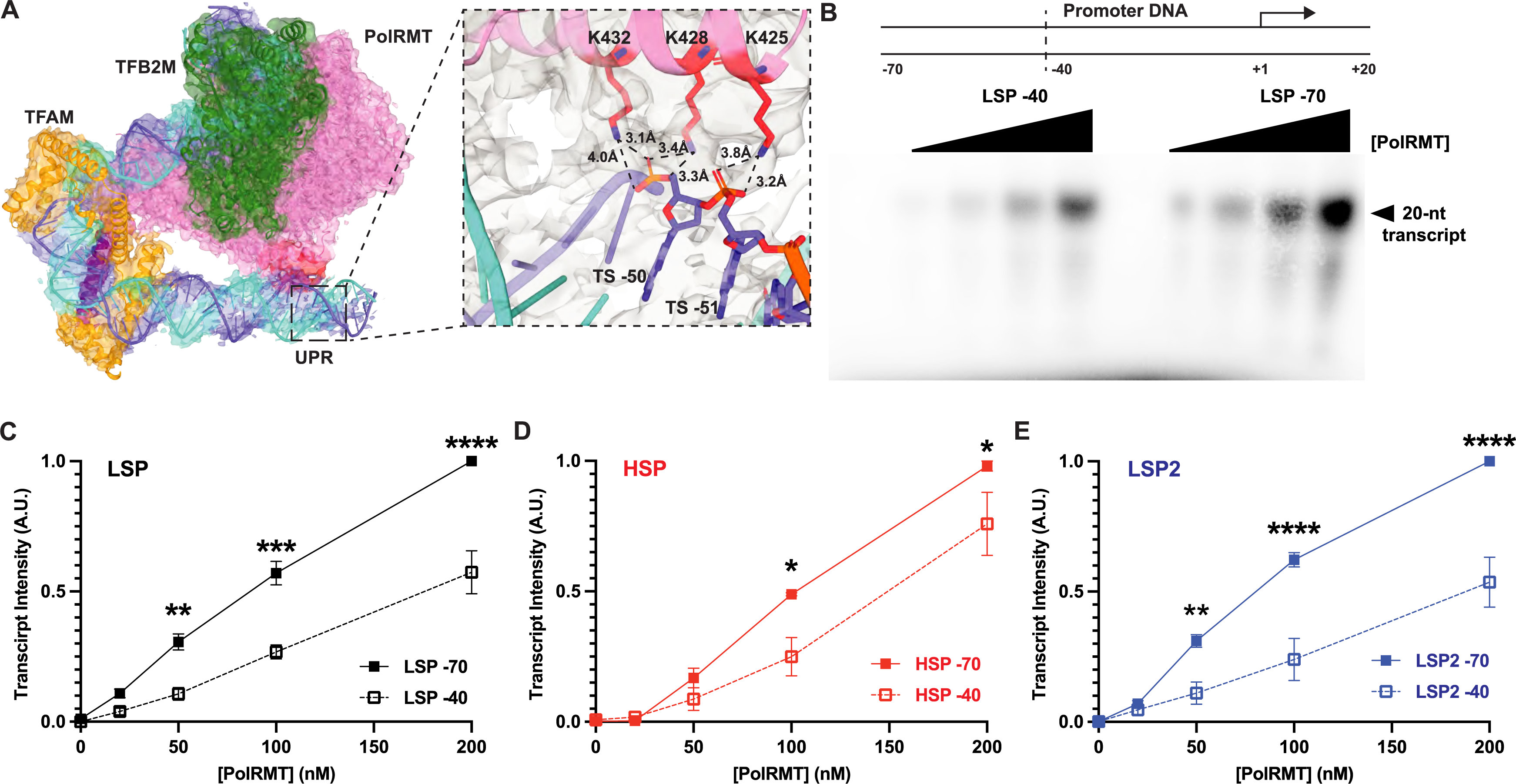
PolRMT interacts with the upstream promoter region (UPR) during transcription initiation. (A) Cryo-EM density map highlighting the interaction of PolRMT K425, K428, and K432 with the UPR, specifically, the DNA backbone of the TS at bases −50 and −51. (B) DNA substrate design for transcription initiation assays with the short template extending to the −40-bp and long template to the −70-bp. Representative gel image of PolRMT titration from 0-200 nM with the −40 and −70-bp LSP DNA template. (C) Quantification of transcript production from the LSP with −40 and −70-bp template. (D) Quantification of transcript production from the HSP with −40 and −70-bp template. (E) Quantification of transcript production from the LSP2 with −40 and −70-bp template. Mean and SEM were determined from 3 independent replicates. Significance was determined using a 2-way ANOVA test with Šídák’s multiple comparisons test. Significance: * p<0.05, ** p<0.01, *** p<0.001, **** p<0.0001.

### Mutation of PolRMT residues K425, K428, and K432 abolishes UPR enhancement of transcript yield

To directly test the role of PolRMT residues K425, K428, and K432 in mediating interaction with the UPR, we mutated all three lysines to glutamate (K425E/K428E/K432E; 3KE) to disrupt electrostatic interactions with the negatively charged DNA backbone. We then evaluated transcription initiation by the 3KE mutant using both long and truncated DNA templates. In contrast to WT PolRMT, transcript yield with the 3KE mutant was largely insensitive to promoter truncation (**Fig. 3A-C**). For LSP, the 3KE mutant produced slightly greater transcript yield from the shorter DNA template, opposite to WT PolRMT (**Fig. 3A**). On both HSP and LSP2, 3KE PolRMT produced similar transcript yield from both templates across all PolRMT concentrations tested (**Fig. 3B** and **3C**). These results confirmed that the interaction between PolRMT and the UPR is mediated primarily by the interaction of PolRMT K425, K428, and K432 with the DNA backbone.

**Figure 3.**
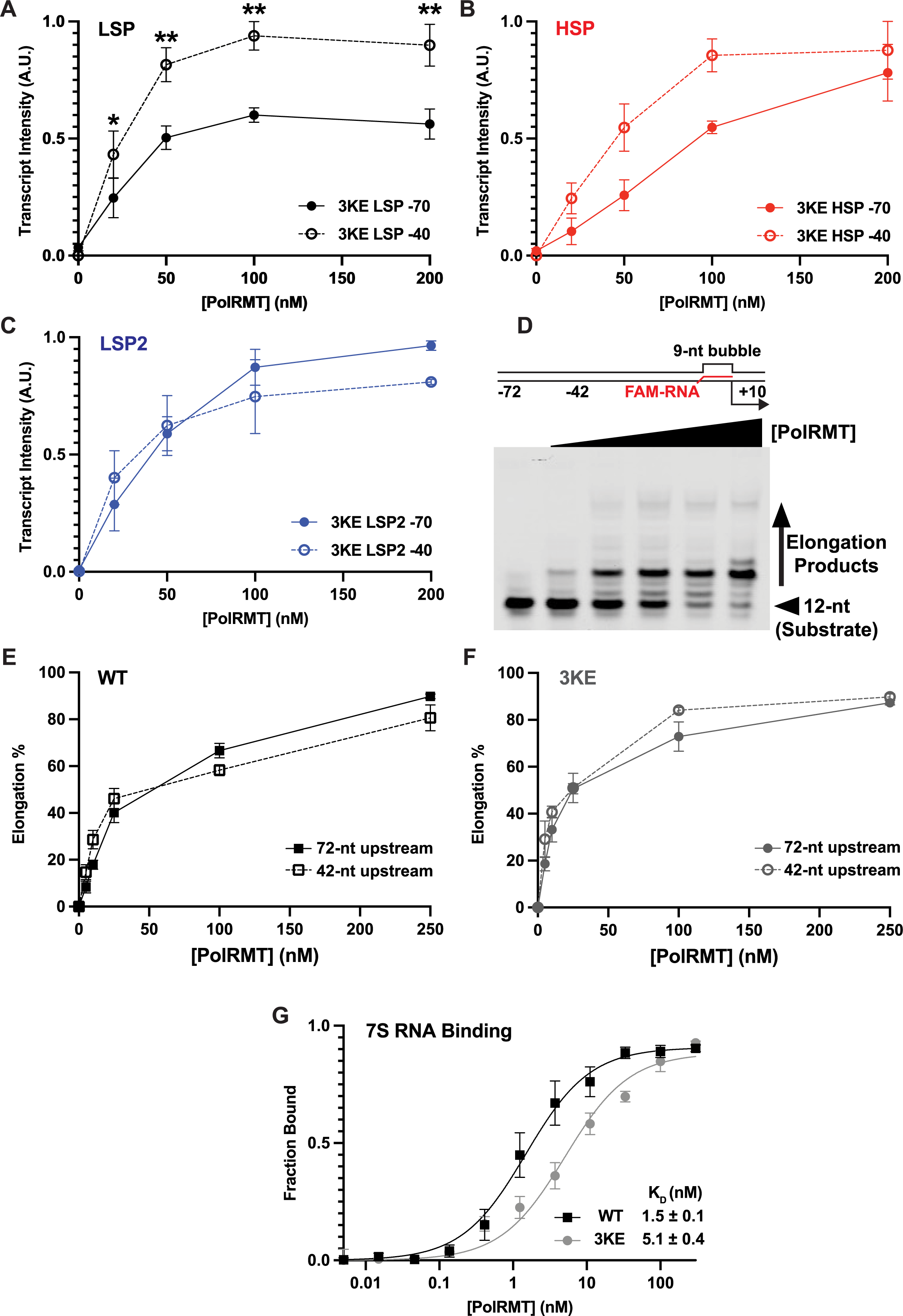
Mutation of PolRMT K425/K428/K432 abolishes the enhancement of transcript yield by the UPR. (A) Quantification of PolRMT K425E/K428E/K432E (3KE) transcript production from the LSP with −40- and −70-bp template. (B) Quantification of PolRMT 3KE transcript production from the HSP with −40- and −70-bp template. (C) Quantification of PolRMT 3KE transcript production from the LSP2 with −40- and −70-bp template. Mean and SEM were determined from 3 independent replicates. Significance was determined using a 2-way ANOVA test with Šídák’s multiple comparisons test. Significance: ** p<0.01. (D) Elongation substrate used for promoter-free transcription elongation assays. The 12-nt FAM-labeled RNA is shown in red and the length of upstream DNA is indicated as either 42-nt or 72-nt. Representative gel of promoter-free transcription elongation assay with titration of PolRMT from 0-250 nM. The unelongated 12-nt substrate and elongation products greater than 12-nt are indicated. (E) Quantification of promoter-free transcription elongation by WT PolRMT on the −72- and −42-nt elongation scaffolds. (F) Quantification of promoter-free transcription elongation by 3KE PolRMT on the −72- and −42-nt elongation scaffolds. Mean and SEM were determined from 3 independent replicates. (G) Quantification of filter binding assays for PolRMT WT and 3KE binding to 7S RNA. Mean and SD were determined from 5 independent replicates.

To test whether this interaction may also facilitate transcription elongation, we employed a promoter-independent transcription elongation assay. The elongation scaffold was designed with a 9-nt bubble between the NT and TS where the labeled RNA was paired with the TS, as in previous studies (32, 33), and an upstream DNA length of either 40 or 72-bp (**Fig. 3D**). The length of upstream DNA had no significant impact on transcription elongation for both WT and the 3KE mutant (**Fig. 3E** and **3F**). Taken together, this suggests that the interaction of PolRMT with extended upstream DNA occurs only during transcription initiation, likely facilitated by TFAM bending of promoter DNA into a U-shape. Previous studies suggested that PolRMT binding to 7S RNA negatively regulates transcription initiation (34). The structure of a transcriptionally-inactive PolRMT dimer bound to 7S RNA suggested that, along with several other positively charged residues, K425, K428, and K432 may contact 7S RNA (34). To probe whether the positively charged residues K425, K428, and K432 may contribute to transcription regulation via RNA interactions, we evaluated binding of PolRMT to 7S RNA using a filter binding assay. We saw a 3-fold decrease in binding affinity in the 3KE mutant compared to WT (**Fig. 3G**). These findings suggest that K425, K428, and K432 contribute not only to transcription initiation but may also contribute to regulation via the extensive PolRMT–7S RNA binding interface.

### Cryo-EM Structure of the Mitochondrial Transcription Initiation Complex without TFAM

During cryo-EM data processing (**Fig. S1** and **Table 1**), we observed a second class of mtTIC in which TFAM was not present and short linear upstream DNA was observed (**Fig. 4A** and **Fig. S3**). This structure was resolved to 2.86 Å resolution, and it lacked density for DNA upstream of position −24 and the PolRMT N-terminal extension (NTE, residues 44-121 and 147-217) (**Fig. 4A**). Notably, we observed electron density adjacent to the short, linear upstream DNA that can accommodate the PolRMT tether helix, which contains several positively charged residues and could interact with the DNA backbone (**Fig. S4**). Aside from the extra density and slightly longer upstream DNA, the core PolRMT and TFB2M of this structure were nearly identical to our structure containing TFAM with an RMSD of 0.6 Å for the 1331 Cα atoms (**Table S2** and **S3**).

**Figure 4.**
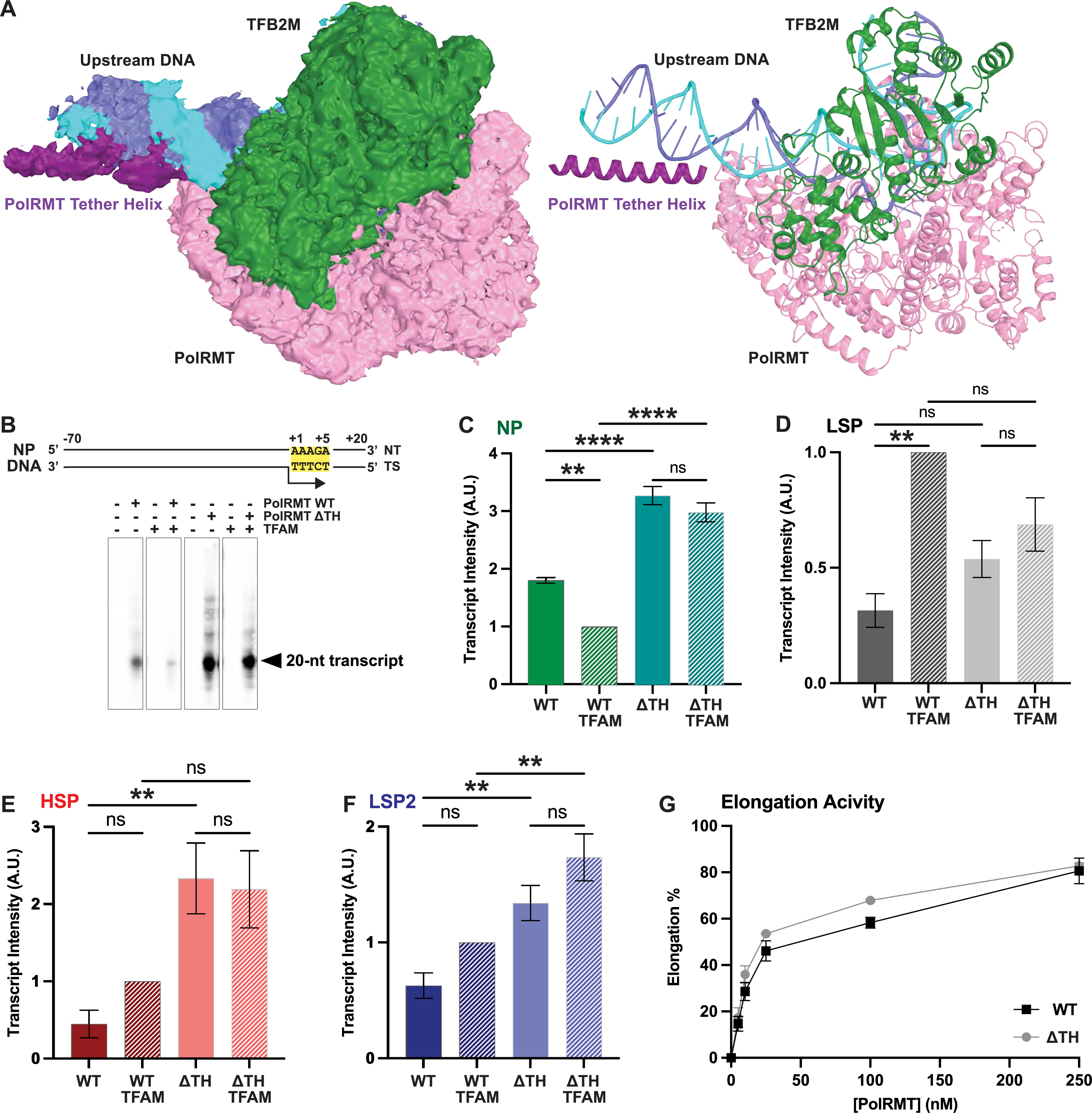
The PolRMT tether helix has an autoinhibitory effect on nonspecific transcription initiation and initiation from the HSP and LSP2. (A) Left: cryo-EM density map of the mtTIC without TFAM, and a short linear upstream DNA with adjacent electron density that accommodates the PolRMT tether helix. Right: Cartoon representation of DNA and proteins colored as in (Fig. 1A). (B) DNA sequence for non-promoter (NP) transcription initiation template that contains the +1 to +5 sequence conserved amongst all three promoters (highlighted yellow) and is derived from the mtDNA genome, positions 4211-4300 (3). Representative gel image of transcription initiation by PolRMT WT or ΔTH in the presence or absence of TFAM on the NP DNA template with the 20-nt transcription product indicated. Transcription reactions contained 20 nM DNA template, 100 nM TFB2M, 0 or 60 nM TFAM, and 0 or 100 nM PolRMT. (C) Quantification of transcript production from the NP DNA substrate. (D) Quantification of transcript production from the LSP. (E) Quantification of transcript intensity from the HSP. (F) Quantification of transcript production from the LSP2. (G) Quantification of promoter-free transcription elongation by WT and ΔTH PolRMT on a −42-nt elongation scaffold. Mean and SEM were determined from 3 independent replicates. Significance was determined using a 1-way ANOVA test with Tukey’s multiple comparisons test. Significance: * p<0.05, ** p<0.01, **** p<0.0001.

### The tether helix of PolRMT influences promoter selection through an autoinhibitory effect

The apparent density for the PolRMT tether helix near the linear upstream DNA suggested a possible role in transcription initiation in the absence of TFAM. To test this, we generated a PolRMT mutant lacking the tether helix (residues 122-146) (ΔTH). Previous work on mouse PolRMT demonstrated that truncation of the entire N-terminal extension (including the tether helix) led to off-target transcription initiation (25). To test non-promoter transcription (NP) initiation, we employed a DNA fragment from the mtDNA genome that contains the conserved AAAGA sequence present at positions +1 to +5 on all three promoters and occurs at over 30 locations throughout the mtDNA genome but lacks the promoter TFAM binding site (**Fig. 4B**). We performed *in vitro* transcription assays to test the transcript yield from WT and ΔTH PolRMT in both the presence and absence of TFAM. The ΔTH mutant had significantly higher NP transcript yield than WT, independent of the presence of TFAM (**Fig. 4C**). Inclusion of TFAM significantly decreased NP transcription of WT PolRMT, while only a slight but not significant reduction was observed for ΔTH (**Fig. 4C**). Further, we used this assay to test each of the three mitochondrial promoters. On LSP, inclusion of TFAM increased transcript yield by WT PolRMT, while ΔTH had similar yield to WT in the absence of TFAM and was not significantly enhanced by the presence of TFAM (**Fig. 4D**). On HSP, ΔTH had significantly greater TFAM-independent transcription yield than WT, while TFAM-dependent transcription yield was slightly higher than WT (**Fig. 4E**). On LSP2, ΔTH had greater transcript production in both the presence and absence of TFAM compared to WT (**Fig. 4F**). A similar trend was also observed when the PolRMT concentration was increased (**Fig. S5**). Additionally, promoter-free transcription elongation was nearly identical for WT and ΔTH PolRMT (**Fig. 4G**). Taken together, these results suggest that the PolRMT tether helix has an autoinhibitory role in transcription initiation from the HSP and LSP2 promoters, and that it contributes to the specificity of transcription initiation by limiting non-promoter initiation.

## Discussion

Human mtDNA transcription initiation is a streamlined, three-component system that relies on the coordinated activities of PolRMT, TFB2M, and TFAM. TFAM is unique among transcription factors in that it also functions as the principal mitochondrial genome-packaging protein (35). Here we identify critical interactions between three positively charged lysine residues of PolRMT and the DNA backbone of the UPR. Our structural and biochemical analyses align well with findings from Morozov et al. (27), suggesting that these interactions are enabled by the TFAM-induced U-shaped DNA architecture promoting productive assembly of the mtTIC. Consistent with this, the UPR enhances transcription initiation on all three mitochondrial promoters, and this stimulatory effect is abolished by mutations of the PolRMT interface that contacts the UPR. Importantly, this interaction is specific to initiation, as elongation proceeds independently of TFAM and is unaffected by these mutations. Notably, the same PolRMT interface also contributes to binding of the transcription-inhibitory 7S RNA (34), as mutations reduce 7S RNA binding, suggesting a shared binding surface that may influence transcription regulation.

The HSP and LSP in the mitochondrial genome are separated by a 153-bp non-coding region. Previous footprinting studies suggested that multiple TFAM molecules may bind within this region to stimulate transcription from both promoters (28). However, in available TIC structures (15, 23, 24), TFAM-induced U-shaped DNA bending positions the immediate upstream DNA in close proximity to the PolRMT surface, such that additional TFAM binding at positions −30 to −60 would sterically clash. Instead, the ∼30-bp region between the two UPRs could accommodate additional TFAM occupancy. Additional TFAM binding at these distal sites may further remodel the non-coding region, potentially bringing the LSP and HSP initiation complexes into spatial proximity (**Fig. 5A**). This could generate a transcriptional hub that coordinates promoter usage and enables regulatory crosstalk, similarly to transcription by yeast RNA Pol II, which forms a dimer of the polymerase-mediator complexes simultaneously on two divergent promoters (36). While PolRMT dimerization via 7S RNA inhibits transcription (34), a different interface was found for dimerization of the apo protein (23), and it is unknown if a higher-order organization of both promoters may involve transcriptionally productive PolRMT dimerization. Defining this higher-order topology will require future structural and biochemical studies aimed at visualizing multi-TFAM assemblies and both LSP and HSP TICs on extended promoter templates.

**Figure 5.**
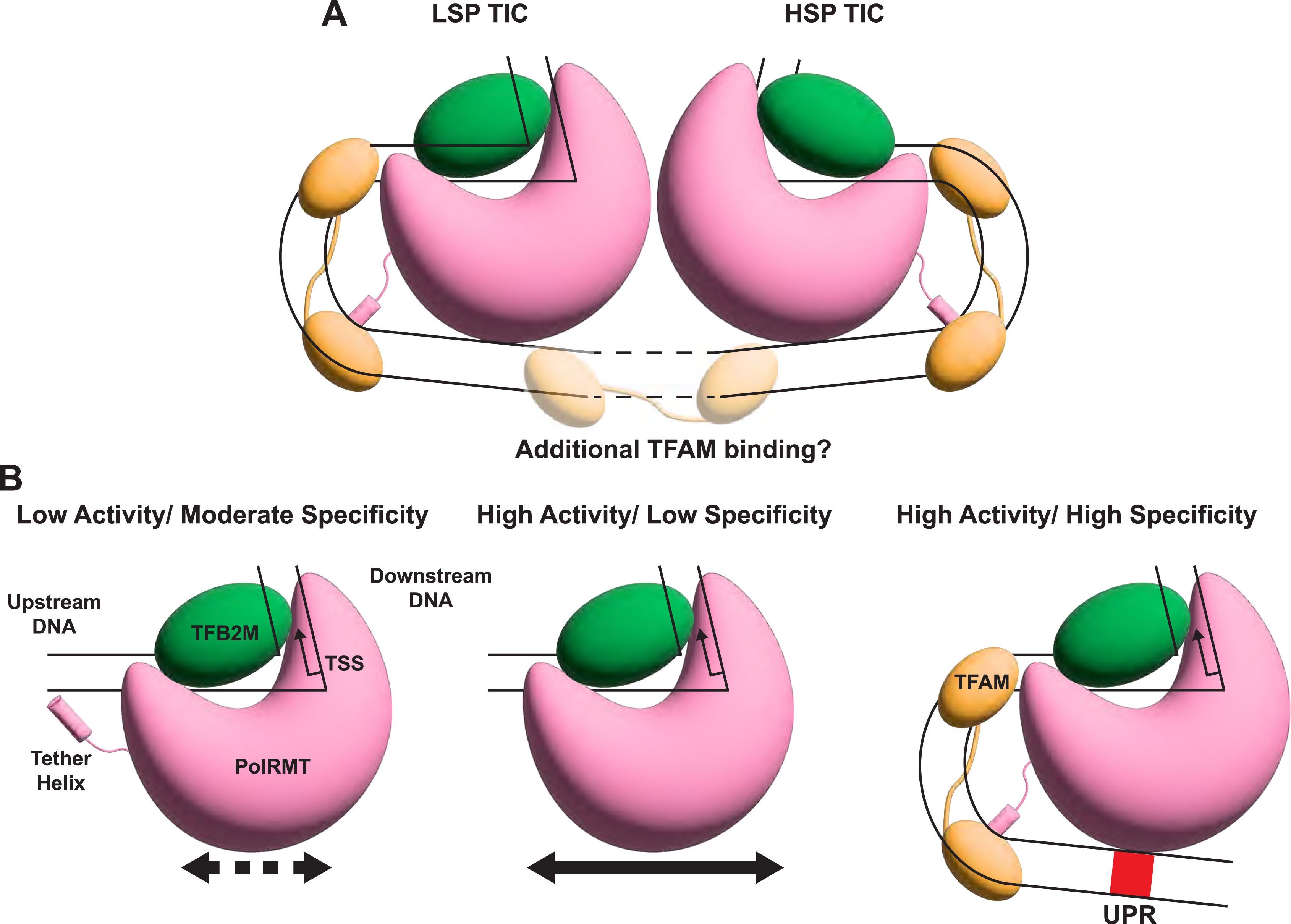
Model for protein–DNA interactions influencing mitochondrial transcription initiation. (A) The mtTIC assembled at both the LSP and HSP ends of the inter-promoter region, with dashed lines to indicate the potential for additional TFAM binding and topological rearrangement. (B) (Left) In the absence of TFAM, PolRMT and TFB2M alone can assemble a mtTIC where the PolRMT tether helix interacts with linear upstream DNA resulting in low activity, likely due to impaired search for the proper TSS, and moderate specificity. (Middle) Truncation of the PolRMT tether helix allows for high transcription activity, possibly due to a faster search for a TSS sequence, but the specificity is poor. (Right) At canonical mtDNA promoters TFAM bends DNA and sequesters the PolRMT tether helix contributing to highly specific transcription initiation while also increasing activity through PolRMT interaction with the UPR.

Another aspect of transcription initiation regulation is the engagement of TFAM and the PolRMT N-terminal tether helix. In the absence of TFAM, we observe the highly positively charged tether helix positioned near the upstream DNA, which remains in a linear conformation. However, this positioning would sterically clash with TFAM-induced U-shaped DNA bending, indicating that the tether helix–DNA interaction is mutually exclusive with productive TFAM binding. Functional assays show that deletion of the tether helix increases non-promoter transcription as well as both TFAM-dependent and TFAM-independent initiation from the HSP and LSP2, supporting an autoinhibitory effect of the tether helix. The nonspecific DNA engagement of the tether helix may inhibit PolRMT activity through impaired ability to scan for the specific TSS. This is analogous to observations of p53 and other DNA-binding proteins that have both specific and nonspecific DNA interactions (37, 38). Alternatively, tether helix–DNA binding may disfavor conformations required for TSS melting (19, 39, 40). Furthermore, the tether helix may adopt intramolecular contacts within PolRMT that reduce nonspecific DNA binding. The presence of TFAM at canonical promoter sites does not allow the tether helix to interact with upstream DNA as it forms electrostatic interactions with TFAM’s acidic surface (**Fig. S4)**, thereby preventing autoinhibition. Although the precise mechanism remains to be defined, our data support a model in which the tether helix suppresses spurious initiation and is repositioned only when proper promoter architecture with TFAM binding is established.

Because TFAM binds DNA with limited sequence specificity and is highly abundant in mitochondrial nucleoids, a key question is how off-target transcription is avoided. Our findings suggest a multilayered mechanism. First, the tether helix suppresses basal PolRMT transcription on non-promoter templates (**Fig. 5B**). Second, productive initiation requires a precise DNA geometry generated by TFAM-induced U-bending and UPR contact with PolRMT. It has been shown that TFAM also contributes to this specificity as the mitochondrial promoters allow for greater DNA bending and complex stability than non-promoter DNA (41, 42). Additionally, high TFAM occupancy of mtDNA directly inhibits PolRMT access and transcription (21). Together, these features establish a coincidence-detection mechanism in which DNA sequence, TFAM-mediated architecture, and polymerase conformational control must align to permit specific and efficient initiation.

Evolutionarily, TFAM was not originally a dedicated transcription factor. The yeast homolog, Abf2p, primarily functions in mitochondrial genome packaging (43–45), and mitochondrial RNA polymerases in these species lack a comparable N-terminal tether helix (**Fig. S6**). This correlation suggests that acquisition of the tether helix and the transcriptional role of TFAM co-evolved to enable tighter control of transcription initiation in metazoan mitochondria. As mitochondrial genomes became more compact and gene expression more tightly coordinated with cellular physiology, TFAM-mediated genome packaging and transcription became functionally integrated, with TFAM coupled to an autoinhibitory polymerase element. This arrangement may have provided an efficient means to enhance promoter specificity within a minimal transcription system.

In summary, our study demonstrates that extended upstream promoter DNA directly engages PolRMT to enhance transcription initiation, while the tether helix can exert an autoinhibitory effect that enforces promoter specificity. By integrating structural and biochemical analyses, we provide a refined model in which TFAM-mediated DNA architecture and PolRMT–DNA interactions cooperate to influence transcription initiation. More broadly, these findings illustrate how modular extensions appended to a bacteriophage-like RNA polymerase can introduce regulatory complexity into a streamlined transcription apparatus. Future studies aimed at defining higher-order promoter DNA organization and regulatory RNA interactions will further clarify how mitochondrial transcription is dynamically tuned under physiological and pathological states.

## Experimental procedures

### Cloning and Expression of TFAM, TFB2M, and PolRMT variants

Human TFAM (residues 43-246) with an N-terminal 6X-His tag and PreScission Protease cleavage sequence was inserted into the pET-28p vector via In-fusion cloning (Takara Bio). Expression was performed in *E. coli* BL21(DE3) and cultures were grown in LB at 37°C until reaching optical density (OD600) of 0.6-0.8. Cultures were cooled on ice for 30 min. Then, expression was induced with 1 mM IPTG, and cultures were incubated in a shaker at 20°C for 18 hours.

Human TFB2M (residues 21-396) was inserted into the pET-SUMO vector with an N-terminal 6X-His-SUMO2 tag via In-fusion cloning. Expression was performed in *E. coli* BL21(DE3) and cultures were grown in LB at 37°C to OD600 = 0.6-0.8. Cultures were cooled on ice for 30 min. Then, expression was induced with 0.5 mM IPTG, and cultures were incubated in a shaker at 16°C for 18 hours.

Human PolRMT WT (residues 44-1230) was inserted into the pMAL-c6T (NEB) vector with an N-terminal 6X-His-MBP tag and PreScission protease cleavage sequence via In-fusion cloning. The vector was transformed into ArcticExpress (Agilent) cells containing the pTF16 plasmid (Takara Bio) expressing TriggerFactor to enhance solubility. Cultures were grown in LB containing 0.5 mg/mL L-Arabinose and 20 g/L glucose at 37°C to OD600 = 0.3. Cultures were cooled on ice for ∼2 hours before inducing expression with 0.1 mM IPTG, then shaking continued at 10°C for ∼40 hours.

The K425E/K428E/K432E (3KE) PolRMT mutant construct was generated from the plasmid containing WT via In-fusion cloning. The tether helix truncation mutant (deletion of PolRMT residues 122-146 [ΔTH]) was generated via inverse PCR and the KLD enzyme mix (NEB) (primers included in **Table S1**).

### Purification of TFAM, TFB2M, and PolRMT variants

TFAM was purified by first resuspending cell pellets in buffer N1 (20 mM Tris pH 7.5, 1 M NaCl, 1 mM BME, 10 mM Imidazole, and 10% glycerol) with 1 mM PMSF, 5 μg/mL DNase, and one Pierce EDTA-free protease inhibitor cocktail tablet. After lysis by sonication, the clarified lysate was incubated for 30 minutes with Ni^2+^ resin (0.5 mL per 1 L cell culture). The mixture was added to a gravity column, washed with buffer N2 (20 mM Tris pH 7.5, 1 M NaCl, and 10% glycerol) containing 30 mM imidazole, and eluted with buffer B containing 300 mM imidazole. The eluent was incubated with PreScission protease at 4°C for 2 hours to cleave the His tag. The solution was diluted to 167 mM NaCl in MS buffer (20 mM Tris pH 7.5 and 3 mM DTT) and loaded onto a Mono S 10/100 GL (Cytiva) column equilibrated in MS buffer with 167 mM NaCl. TFAM was eluted with a gradient of MS buffer containing 1 M NaCl, with peak elution near 500 mM NaCl. Peak fractions were concentrated, aliquoted, and stored at −80°C.

TFB2M was first purified by nickel affinity as described for TFAM. The eluent was then incubated with SenP2 at 4°C for 2 hours to cleave the His-SUMO tag. The sample was diluted to 250 mM NaCl in Hep buffer (40 mM Tris pH 7.5 and 3 mM DTT) and loaded onto a 5 mL HiTrap Heparin HP (Cytiva) column equilibrated in Hep buffer containing 250 mM NaCl. TFB2M was eluted with a gradient of Hep buffer containing 1 M NaCl, with peak elution near 500 mM NaCl. Peak fractions were concentrated, aliquoted, and stored at −80°C.

PolRMT WT, 3KE, and ΔTH were purified by resuspending cells in buffer M1 (1X PBS, 1 M NaCl, 20% glycerol, 5 mM EDTA, 3 mM DTT, and 1 mM PMSF) containing 5 μg/mL DNase and 1 Pierce EDTA-free protease inhibitor cocktail tablet. Cells were lysed by sonication, and the clarified lysate was incubated for 1.5 hours with Amylose resin (250 μL per 1 L cell culture). The mixture was loaded onto a gravity column, washed with buffer M2 (40 mM Tris pH 8.0, 1 M NaCl, 20% glycerol, 1 mM EDTA, 5 mM DTT, and 1 mM PMSF), and eluted by incubating overnight with buffer M3 (40 mM Tris pH 8.0, 1 M NaCl, 20% glycerol, 1 mM EDTA, 5 mM DTT, 1 mM PMSF, and 300 mM Maltose). The eluent was diluted to 250 mM NaCl in QH buffer (40 mM Tris pH 8.0, 5 mM DTT, 1 mM EDTA, and 10% glycerol) and loaded onto two columns connected in series: 1 mL HiTrap Capto Q column (Cytiva) followed by 5 mL HiTrap Heparin HP column. The columns were equilibrated in QH buffer containing 250 mM NaCl. After loading, the Capto Q column, which contained contaminating nucleic acid, was removed prior to elution. Finally, the heparin column was eluted with a gradient of QH buffer with 1 M NaCl, with peak elution near 400 mM NaCl. Peak fractions were concentrated, aliquoted, and stored at −80°C.

### *In Vitro* Transcription Initiation Assay

DNA templates for *in vitro* transcription assays were generated by annealing complementary oligonucleotides (IDT) (**Table S1**). Transcription initiation reactions contained 20 nM DNA, 100 nM TFB2M, 60 nM TFAM, and 0-300 nM PolRMT variant (presence of TFAM and PolRMT concentrations are indicated in figures) in transcription buffer (40 mM Tris pH 8.0, 10 mM MgCl_2_, 100 μg/mL BSA, 1 mM DTT, and 2 U/μL RNase inhibitor, Murine [NEB]) with 500 μM ATP/CTP/GTP, 10 μM cold UTP, and 33.3 nM ^32^P α-UTP. Reactions were incubated for 30 min at 30°C and stopped by the addition of 2X Urea-PAGE loading buffer (90% formamide, 50 mM EDTA, 0.1% xylene cyanol, and 0.1% bromophenol blue) followed by incubation at 95°C for 10 min. Reactions were resolved by UREA-PAGE on a 20% polyacrylamide gel in 1X TBE buffer with 30 min pre-run followed by a 1.5-hour run at 250V. Gels were exposed to phosphor imaging screen overnight, scanned, and quantified using a Sapphire Biomolecular Imager and Azure Spot (Azure Biosystems).

### Promoter-Free *In Vitro* Transcription Elongation Assay

Elongation scaffolds for *in vitro* transcription elongation assays were generated by annealing DNA oligonucleotide pairs with a 5’-FAM-labelled RNA oligonucleotide (IDT) (**Table S1**). Annealing was performed by mixing the non-template strand, template strand, and RNA in a 2:1.5:1 molar ratio, heating to 90°C for 5 min, cooling to 60°C at a rate of −0.5°C per min, holding at 60°C for 20 min, followed by cooling to 12°C at −0.5°C per min. Transcription elongation reactions contained 20 nM annealed elongation scaffold and 0-250 nM PolRMT variant (concentrations are indicated in figures) in transcription buffer with 500 μM ATP/CTP/GTP/UTP. Reactions were incubated for 20 min at 30°C and stopped by the addition of 2X Urea-PAGE loading buffer followed by incubation at 95°C for 10 min. Reactions were resolved by UREA-PAGE on a 22.5% polyacrylamide gel in 1X TBE buffer with 30 min pre-run followed by a 1.5-hour run at 250V. Gels were scanned and quantified using a Sapphire Biomolecular Imager and Azure Spot.

### Double Filter Binding Assay

The 7S RNA substrate for binding assays was *in vitro* transcribed using T7 RNAP, dephosphorylated with calf intestinal phosphatase (NEB), 5’ end-labeled with [γ-^32^P] ATP (Revvity) using T4 polynucleotide kinase (NEB), and purified by denaturing PAGE. 0.1 nM radiolabeled 7S RNA was incubated with PolRMT (0-300 nM) in binding buffer (40 mM HEPES pH 7.5, 100 mM KCl, 20 mM β-ME) at 25°C for 15 minutes. Reactions were vacuum-filtered through nitrocellulose (0.45 μm, Schleicher & Schuell) and nylon (Whatman) membranes in a dot-blot microfiltration apparatus (Bio-Rad) (46). Membranes were air-dried and quantified by phosphorimaging (Cytiva). The fraction of RNA bound was calculated by the ratio of counts on nitrocellulose to total counts (nitrocellulose + nylon) (47).

### Cryo-EM Sample Preparation

The complex was assembled by mixing 3.3 μM TFAM, 5 μM TFB2M, 3.3 μM PolRMT WT, and 3.6 μM HSP −60 to +11 DNA substrate yielding a final concentration for the complex of 1 mg/mL in buffer containing 40 mM Tris (pH 8.0), 10 mM MgCl_2_, 10 mM DTT, and 100 mM NaCl. After 30 min incubation on ice and centrifugation at 17,000 x *g* for 10 min, a 3 μL drop of the complex was spotted onto freshly glow-discharged grids (Quantifoil R 1.2/1.3 Cu 300 mesh grids). Excess sample was blotted using the Vitrobot Mark IV (FEI) with the standard Vitrobot filter paper (Ø55/20 mm [Ted Pella]), with blotting time 2 s and blotting force 3 under 100% humidity at 20°C. The grids were flash-frozen in liquid ethane and stored in liquid nitrogen.

### Cryo-EM Data Collection and Processing

The dataset of 27,688 movies was collected from Stanford-SLAC Cryo-EM Center and recorded on a Titan Krios G4i electron microscope operated at 300 kV with serial EM (48) (detailed parameters for data collection are summarized in **Table 1**). Motion correction was performed with MotionCor2 (49), and defocus values were estimated on non-dose-weighted micrographs with Gctf (50). For processing of the dataset, reference-free auto-picking (Laplacian-of-Gaussian picking) was performed in RELION-4.0 (51, 52). 11,635,618 particles were picked and extracted to pixel size 4.79 Å/pixel and imported to cryoSPARC-4.2 for 2D classification for multiple rounds of 2D classification and subset selection. 7,872,349 particles were used for ab-initio reconstruction and separated into six classes. Four of those classes were subjected to heterogeneous refinement, where one class showed clear features for PolRMT, TFB2M, and DNA. Particles from this class were further cleaned with 2D classification and split into two classes by ab-initio reconstruction and heterogeneous refinement. The first class was further processed to yield the HSP without TFAM complex, while the second class was used for the HSP with TFAM complex. Particles from the first class with only density for short upstream DNA were extracted (0.958 Å/pixel) in RELION-4.0 and reimported to cryoSPARC-4.2 for ab-initio reconstruction into six classes followed by heterogeneous refinement and non-uniform refinement. Particles from the three classes with density for DNA were subjected to heterogeneous refinement with the best map (3.37 Å resolution) and worst map from the previous non-uniform refinement. This yielded 446,280 particles, which were used for CTF refinement and non-uniform refinement resulting in a map with a resolution of 2.86 Å. Particles from the class with density for longer DNA were extracted (0.958 Å/pixel) in RELION-4.0 and reimported to cryoSPARC-4.2 for ab-initio reconstruction into six classes followed by heterogeneous refinement and non-uniform refinement. One class yielded a 3.51 Å resolution map with density for the extended DNA, which was used as a template for Topaz (53) training and particle picking. In parallel, this class was subjected to local 3D classification using a focused mask for the extended DNA, which resulted in 2 of 10 classes with density for the DNA in contact with PolRMT. Non-uniform refinement of these two classes produced a 4.13 Å resolution map. Topaz particle picking resulted in 32,940,939 particles which were extracted (0.958 Å/pixel) in RELION-4.0 and reimported to cryoSPARC-4.2 for 2D cleaning resulting in 23,147,478 particles. The particles were then subjected to heterogeneous refinement with the 4.13 Å resolution map combined with five poor maps from previous ab-initio reconstruction. This produced one class with features of PolRMT, TFB2M, TFAM and extended DNA, which was used for three rounds of iterative ab-initio reconstruction and heterogeneous refinement. This yielded 200,388 particles, which were used for CTF refinement and non-uniform refinement resulting in a map with a resolution of 3.33 Å. Both maps were subject to 3DFlex jobs (54) to improve local resolution and Phenix auto sharpen (55) to aid in model building.

### Model Building and Refinement

For the HSP with TFAM complex, the previously published structure of the TIC HSP crystal structure (PDB: 6ERQ) was used as an initial model for manually docking into the cryo-EM density map using UCSF Chimera (56). Additionally, an AlphaFold 3 (57) prediction of the mtTIC with the extended HSP substrate was used for modelling the DNA, and the recently published structures of the pre-IC3 (PDB 9GZM) and IC0 (9MN5) on LSP were used for modeling regions of the PolRMT NTE, thumb, and specificity loop and TFB2M NTD. The model was further manually rebuilt in COOT (58) based on electron density and refined in Phenix (59) with real-space refinement and secondary structure and geometry restraints. For the HSP without TFAM complex, the model of the HSP with TFAM complex was used as an initial model for manually docking into the cryo-EM density map using Chimera. An AlphaFold 3 prediction of the mtTIC without TFAM was used to model the linear upstream DNA. As before, the model was manually rebuilt in COOT and refined in Phenix.

### Statistical Analysis

All data were graphed and analyzed for significance using GraphPad Prism 10. For transcription initiation reactions with PolRMT titration in **Figure 2**-**3**, background was subtracted from intensity values and normalization was performed by dividing each intensity by the maximum intensity value from each individual experiment gel. Significance was determined using a 2-way ANOVA test with Šídák’s multiple comparisons test. For transcription initiation reactions with a single concentration of PolRMT in **Figure 4** and **Figure S5**, background was subtracted from raw values and normalization was performed by dividing each intensity by the intensity value of the sample containing TFAM and WT PolRMT from each individual experiment gel. Significance was determined using a 1-way ANOVA test with Tukey’s multiple comparisons test. For transcription elongation reactions, elongation percentage was calculated by dividing the intensity of the elongation product bands by the total intensity of the lane after subtracting background. Mean and SEM values were determined from 3 independent replicates for all transcription reactions. For filter binding assays, binding measurements were fit to a one-site specific binding model to determine K_D_ values based on 5 independent replicates. To compare the K_D_ values an extra sum-of-squares F test was performed and the null hypothesis was rejected for p-value < 0.05.

## Protein Sequence Alignments

Sequence alignments of PolRMT homologs and TFAM homologs for **Figure S6** were performed with constraint-based multiple alignment tool (COBALT) (60).

## Data availability

All original data and materials are available upon request.

## Supporting information

This article contains supporting information.

## Acknowledgements

We thank Dr. Gaya P. Yadav at the Laboratory for Biomolecular Structure and Dynamics (LBSD) of Texas A&M University and the core facility at Stanford-SLAC Cryo-EM Center for cryoEM data collection.

## Author contributions

RES purified recombinant proteins, performed *in vitro* transcription assays, cryo-EM sample preparation, data processing, model building, refinement, and statistical analysis, wrote and edited manuscript. CS generated the 7S RNA substrate and performed filter binding assays. XD performed preliminary cryo-EM data processing. JS performed cloning for TFAM and TFB2M. AJH reviewed and edited manuscript. YG conceptualized and administered project, wrote and edited manuscript.

## Funding and additional information

The work was supported by the National Institutes of Health [35GM142722 to Y.G.], and Tufts Faculty Startup Research Fund to AJH and the Cancer Prevention & Research Institute of Texas (CPRIT) [RR190046].

The content is solely the responsibility of the authors and does not necessarily represent the official views of the National Institutes of Health.

## Conflict of interest

The authors declare that they have no conflicts of interest with the contents of this article.

## Abbreviations and nomenclature

AT-RRL: AT rich recognition loop
CTD: C-terminal domain
HMG: high mobility group
HSP: heavy strand promoter
LSP: light strand promoter
LSP2: light stand promoter 2
mtDNA: mitochondrial DNA
MTS: mitochondrial targeting sequence
mtTIC: mitochondrial transcription initiation complex
NP: non-promoter
NT: non-template strand
NTD: N-terminal domain
NTE: N-terminal extension
OXPHOS: oxidative phosphorylation
PolRMT: mitochondrial RNA polymerase
PPR: pentatricopeptide repeat
TFAM: mitochondrial transcription factor A
TFB2M: mitochondrial transcription factor B2
TS: template strand
TSS: transcription start site
UPR: upstream promoter region

## Notes

### Competing Interest Statement

The authors have declared no competing interest.

### Summary of Updates

The manuscript has been revised to change the overall format as well as the content of the abstract, introduction, and discussion. The PDB and EMDB ID has also been removed while we continue to examine the structures.

### Funding

- NIH Common Fund: R35GM142722
- Tufts University

