## Supplementary figures for "Cryo-EM Structures Reveal Upstream DNA Interactions within the Mitochondrial Transcription Initiation Complex"

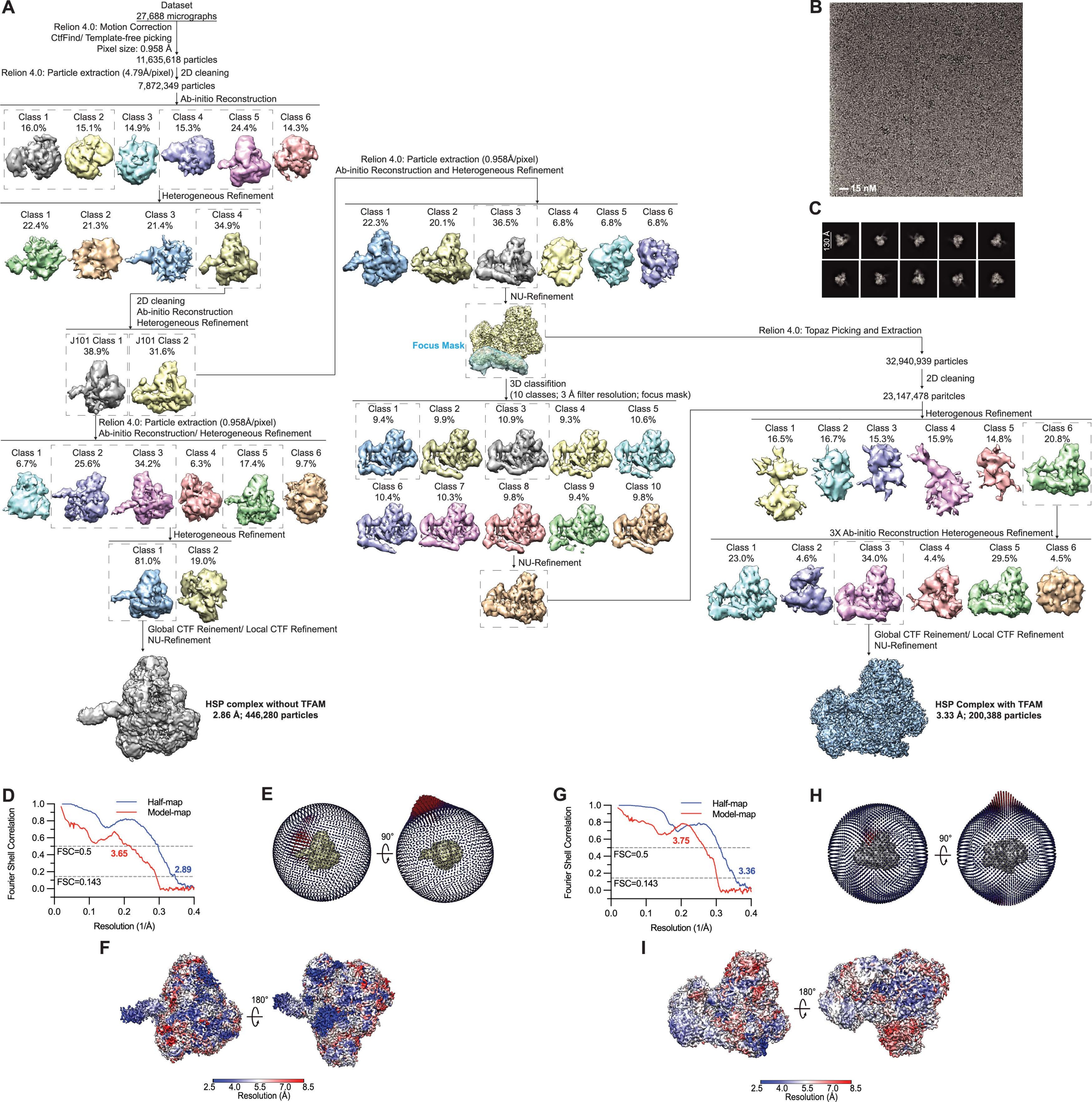
A
Dataset
27,688 micrographs
Relion 4.0: Motion Correction
CtfFind/ Template-free picking
Pixel size: 0.958 Å
11,635,618 particles
Relion 4.0: Particle extraction (4.79Å/pixel)
2D cleaning
7,872,349 particles
Ab-initio Reconstruction
Class 1 16.0%
Class 2 15.1%
Class 3 14.9%
Class 4 15.3%
Class 5 24.4%
Class 6 14.3%
Heterogeneous Refinement
Class 1 22.4%
Class 2 21.3%
Class 3 21.4%
Class 4 34.9%
2D cleaning
Ab-initio Reconstruction
Heterogeneous Refinement
J101 Class 1 38.9%
J101 Class 2 31.6%
Relion 4.0: Particle extraction (0.958Å/pixel)
Ab-initio Reconstruction/ Heterogeneous Refinement
Class 1 6.7%
Class 2 25.6%
Class 3 34.2%
Class 4 6.3%
Class 5 17.4%
Class 6 9.7%
Heterogeneous Refinement
Class 1 81.0%
Class 2 19.0%
Global CTF Reinement/ Local CTF Refinement
NU-Refinement
HSP complex without TFAM
2.86 Å; 446,280 particles
Relion 4.0: Particle extraction (0.958Å/pixel)
Ab-initio Reconstruction and Heterogeneous Refinement
Class 1 22.3%
Class 2 20.1%
Class 3 36.5%
Class 4 6.8%
Class 5 6.8%
Class 6 6.8%
NU-Refinement
Focus Mask
3D classifition
(10 classes; 3 Å filter resolution; focus mask)
Class 1 9.4%
Class 2 9.9%
Class 3 10.9%
Class 4 9.3%
Class 5 10.6%
Class 6 10.4%
Class 7 10.3%
Class 8 9.8%
Class 9 9.4%
Class 10 9.8%
NU-Refinement
Relion 4.0: Topaz Picking and Extraction
32,940,939 particles
2D cleaning
23,147,478 paritcles
Heterogenous Refinement
Class 1 16.5%
Class 2 16.7%
Class 3 15.3%
Class 4 15.9%
Class 5 14.8%
Class 6 20.8%
3X Ab-initio Reconstruction Heterogeneous Refinement
Class 1 23.0%
Class 2 4.6%
Class 3 34.0%
Class 4 4.4%
Class 5 29.5%
Class 6 4.5%
Global CTF Reinement/ Local CTF Refinement
NU-Refinement
HSP Complex with TFAM
3.33 Å; 200,388 particles
B
15 nM
C
130 Å
D
Fourier Shell Correlation
Half-map
Model-map
FSC=0.5
FSC=0.143
3.65
2.89
Resolution (1/Å)
E
90°
G
Fourier Shell Correlation
Half-map
Model-map
FSC=0.5
FSC=0.143
3.75
3.36
Resolution (1/Å)
H
90°
F
180°
2.5 4.0 5.5 7.0 8.5
Resolution (Å)
I
180°
2.5 4.0 5.5 7.0 8.5
Resolution (Å)

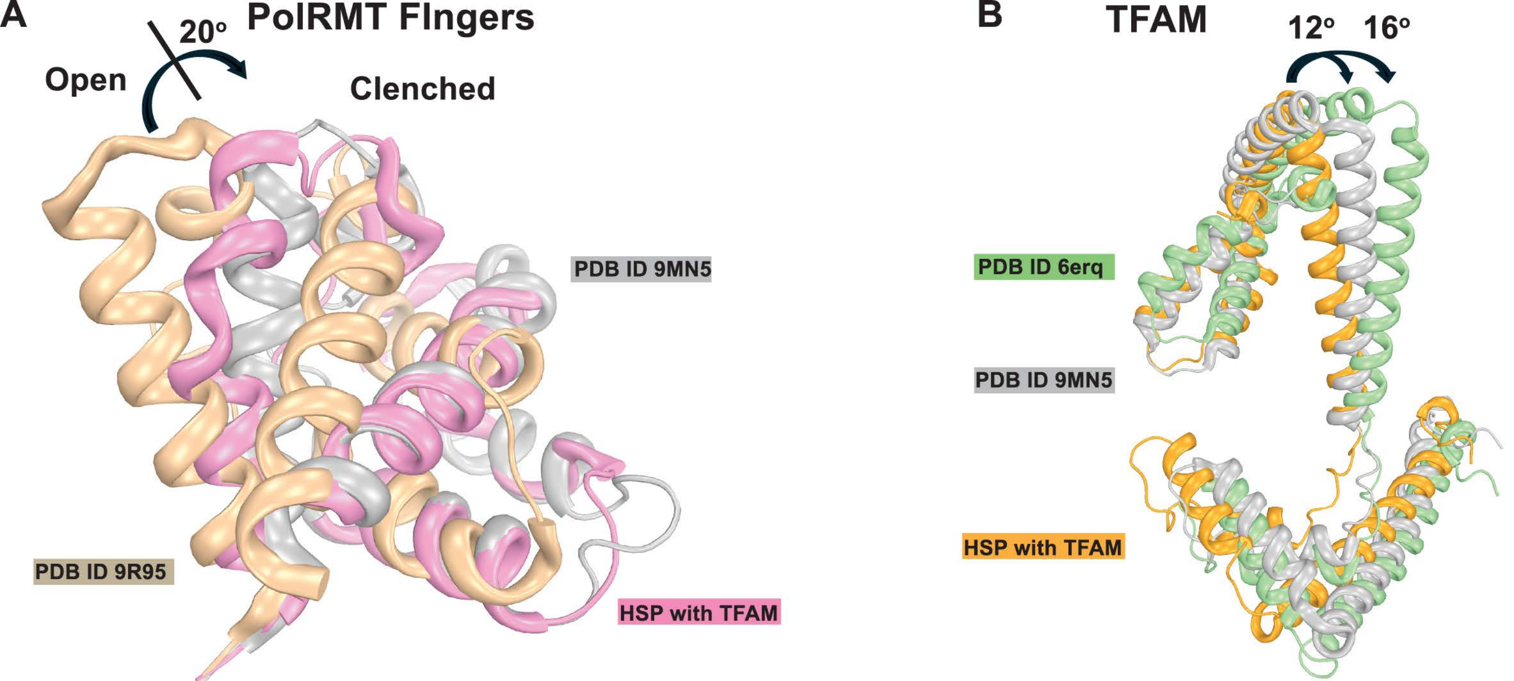
A
PolRMT FIngers
20°
Open
Clenched
PDB ID 9MN5
PDB ID 9R95
HSP with TFAM
B
TFAM
12°
16°
PDB ID 6erq
PDB ID 9MN5
HSP with TFAM

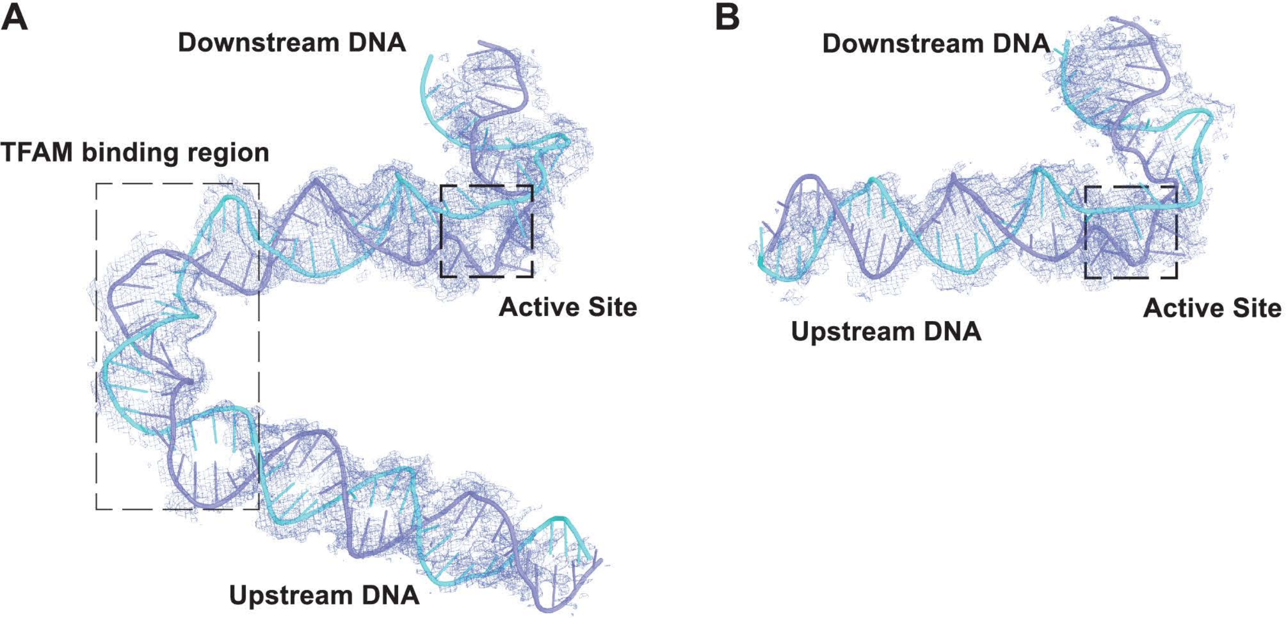
A
Downstream DNA
TFAM binding region
Active Site
Upstream DNA
B
Downstream DNA
Active Site
Upstream DNA

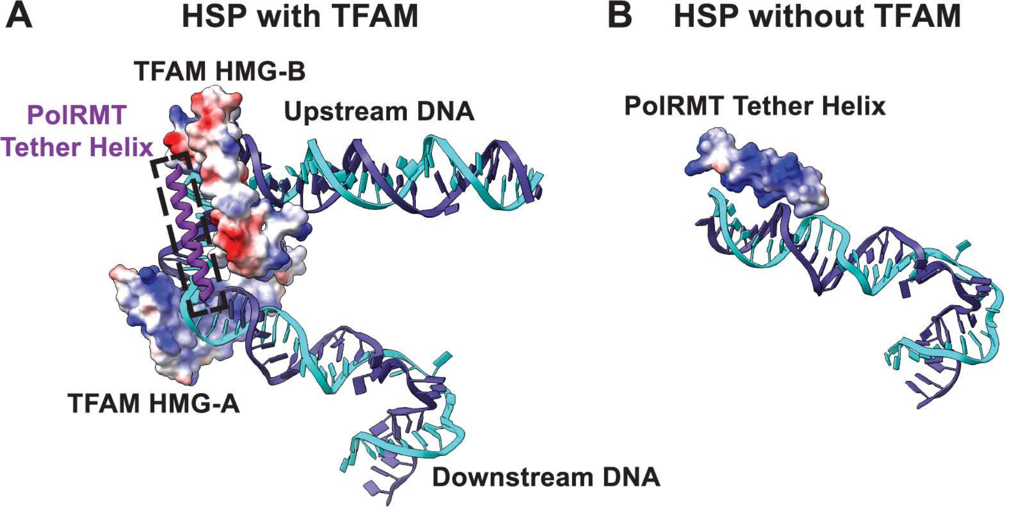
A
HSP with TFAM
TFAM HMG-B
PolRMT
Tether Helix
Upstream DNA
TFAM HMG-A
Downstream DNA
B
HSP without TFAM
PolRMT Tether Helix

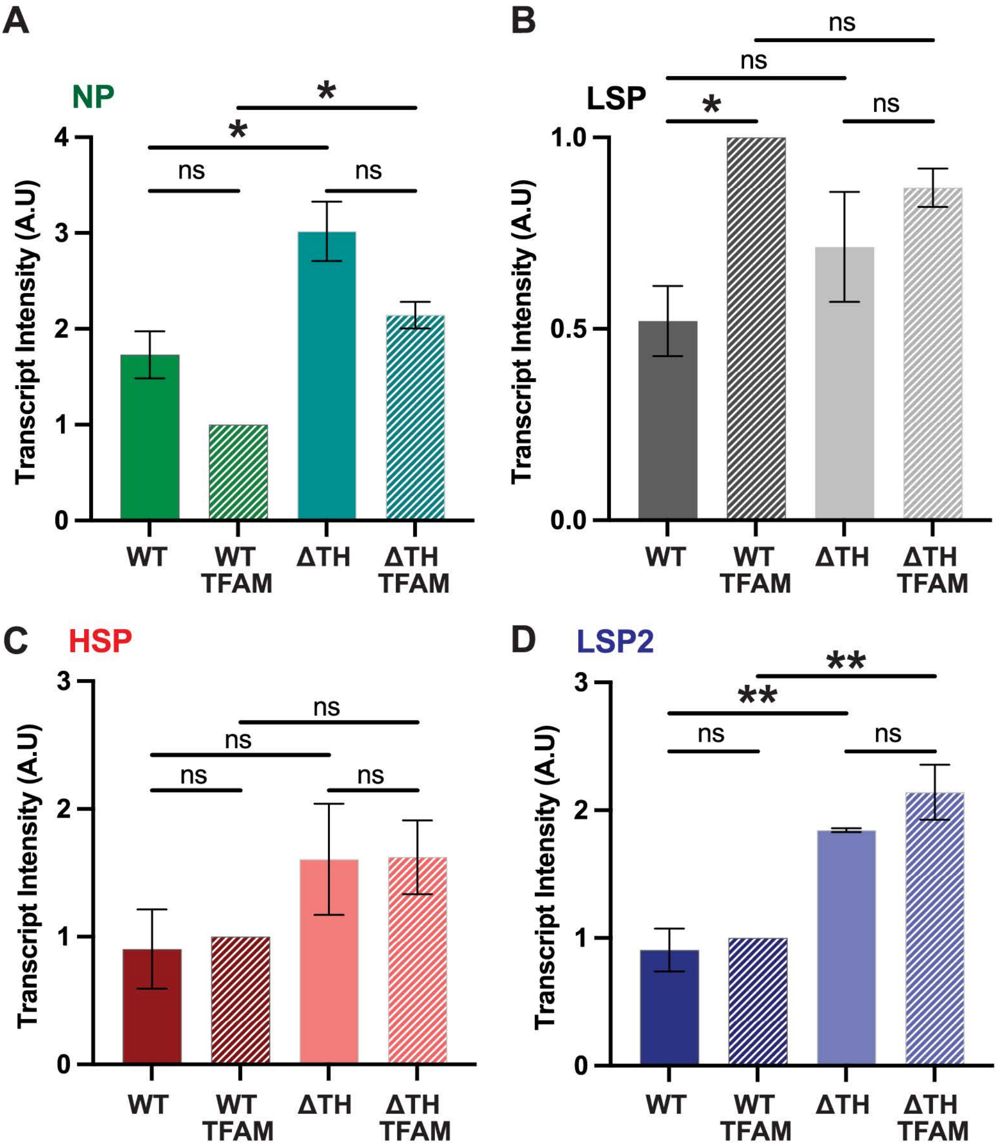
A
NP
Transcript Intensity (A.U)
0
1
2
3
4
ns
*
*
ns
WT
WT TFAM
ΔTH
ΔTH TFAM
B
LSP
Transcript Intensity (A.U)
0.0
0.5
1.0
ns
ns
*
ns
WT
WT TFAM
ΔTH
ΔTH TFAM
C
HSP
Transcript Intensity (A.U)
0
1
2
3
ns
ns
ns
ns
WT
WT TFAM
ΔTH
ΔTH TFAM
D
LSP2
Transcript Intensity (A.U)
0
1
2
3
**
**
ns
ns
WT
WT TFAM
ΔTH
ΔTH TFAM

A

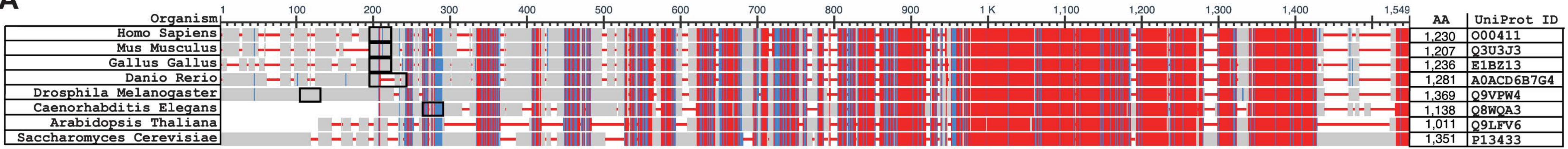


B

| Organism | Potential Tether Helix Sequence | Position |
|---|---|---|
| Homo Sapiens | WAKILEKDKRTQQMRMQRLKAKLQM | 122-146 |
| Mus Musculus | WAQKLEAEKRVKQRRQKEVDQQKQA | 97-121 |
| Gallus Gallus | WTEKLKKEMYIRQLKVEKKLSIAAS | 108-132 |
| Danio Rerio | WMAKLNREMWINPKTKTSKKVIKTG | 156-180 |
| Drosphila Melanogaster | VELLEVTEQAISERRSRVKRLKSKK | 104-128* |
| Caenorhabditis Elegans | KQKKRGIWRQRVKQIALADIVRFTL | 59-83* |
| Arabidopsis Thaliana | N/A | |
| Saccharomyces Cerevisiae | N/A | |

C

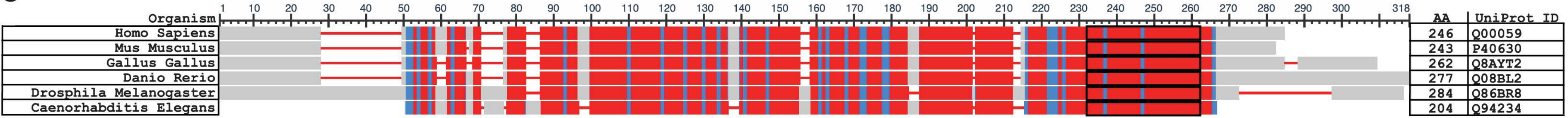


D

| Organism | TFAM HMG-B Helix 3 |
|---|---|
| Homo Sapiens | DSEKELYIQHAKEDETRYHNEMKSWEEQMIE |
| Mus Musculus | PEEKQAYIQLAKDDRIRYDNEMKSWEEQMAE |
| Gallus Gallus | SSQKQPYLQLAQDDKVRYQNEMKSWEAKMVE |
| Danio Rerio | DTQKQMYIQLAEDDKVRYKNEIKSWEEHMME |
| Drosphila Melanogaster | DSEKEVYMQESRKEMELYRKAISVWEEKMIR |
| Caenorhabditis Elegans | DSQKKKYTDEAKKLKDEYHVVLQKWEAEQKE |
